# α2δ-2 mediates coupling of presynaptic calcium entry to vesicle release in hippocampal parvalbumin-expressing interneurons

**DOI:** 10.64898/2026.08.31.748347

**Authors:** Allison J. Ellingson, Emma C. Jerome, Ashlynn A. Gallagher, Eric Schnell

**Affiliations:** Department of Anesthesiology and Perioperative Medicine, Oregon Health & Science University, Portland, OR, 97239; Research and Development Service, Portland VA Health Care System, Portland, OR, 97239, Portland, OR, 97239; Neuroscience Graduate Program, Oregon Health & Science University, Portland, OR, 97239

## Abstract

The α2δ family of auxiliary voltage-gated calcium channel (VGCC) subunits have critical but incompletely understood roles in brain function. Parvalbumin-positive (PV+) interneurons in the hippocampus highly express the α2δ-2 isoform, and mice lacking α2δ-2 exhibit spontaneous seizures. Thus, we examined PV+ neuron-mediated synaptic inhibition in acutely prepared brain slices from *α2δ-2* knockout (KO) mice. In the inner molecular layer of the dentate gyrus, *α2δ-2* KO mice demonstrated an increase in the excitation/inhibition ratio of synaptic inputs onto granule cells. We then used optogenetics to activate PV+ interneurons, which produced dramatically smaller inhibitory synaptic currents in granule cells from *α2δ-2* KO mice. There was a reduction in PV+ inputs onto granule cells as determined by immunostaining. Functionally, these inputs had a lower probability of GABA release and a decreased readily releasable pool of vesicles compared to littermate controls. VGCC coupling to presynaptic vesicle release was also reduced in dentate gyrus PV+ cells in *α2δ-2* KO mice, based on manipulations of intracellular and extracellular calcium. Together, our data indicate that α2δ-2 plays a critical role in PV+ interneuron-mediated synaptic inhibition, which may contribute to seizures in *α2δ-2* mutant mice.

**Significance Statement:** A family of auxiliary VGCC subunits, the α2δs, plays a variety of important roles in brain function, although its underlying mechanisms are poorly understood. Here, we use mouse models to determine that the α2δ-2 isoform plays a critical role in the presynaptic function of parvalbumin-expressing interneurons in the hippocampal dentate gyrus. Using electrophysiological recordings from genetically modified mice, we find that α2δ-2 functionally couples calcium entry to vesicle release from parvalbumin-expressing cells. This finding not only provides a potential explanation for the seizure phenotype of *α2δ-2* mutant mice but also illuminates the broader roles of α2δ proteins in neuronal function.

## Introduction

Voltage-gated calcium channels (VGCCs) critically mediate fast synaptic neurotransmission in the nervous system, and their dysfunction underlies various neurological disorders including epilepsy (Catterall, 2011; Noebels and Jasper, 2012; Nanou and Catterall, 2018). VGCCs consist of a pore forming subunit (α1) and auxiliary subunits (α2δ, β, and γ), which modulate channel properties and subcellular trafficking (Dolphin, 2012). α2δ subunits consist of an extracellular α2 protein linked via a disulfide bond to a δ protein, which tethers this subunit to the cell membrane via a glycophosphatidylinositol (GPI) anchor (Davies et al., 2010). α2δ proteins are of clinical interest as the primary targets for the frequently used medication gabapentin, and because mutations in the genes coding for these subunits cause neurological disease, including epilepsy, autism, and ataxia (Gee et al., 1996; Marais et al., 2001; Iossifov et al., 2012; Pippucci et al., 2013; Punetha et al., 2019).

Although α2δs play minor roles in regulating VGCC gating (Barclay et al., 2001), they influence calcium-dependent neuronal functions in other ways. For example, they modulate calcium current density, likely by trafficking VGCCs to the cell membrane (Shistik et al., 1995; Barclay et al., 2001; Hoppa et al., 2012; D’Arco et al., 2015).

Increasing evidence also suggests that α2δ proteins may also functionally couple calcium entry to downstream effector mechanisms. In hippocampal pyramidal cell cultures, α2δ-1 couples calcium entry to presynaptic vesicle release (Hoppa et al., 2012), and in cerebellar Purkinje cells, α2δ-2 functionally couples postsynaptic VGCCs to endocannabinoid release (Beeson et al., 2022). It is unclear, however, whether the roles of these isoforms in functional coupling are limited to a distinct population of specific sites, or whether they represent a broader role for α2δ proteins in neuronal function across the brain.

Animal studies of α2δs have been challenging due to functional compensation between the α2δ-1 and -3 isoforms (Geisler et al., 2021; Schöpf et al., 2021). Unlike *α2δ-1* or *α2δ-3* KO mice, however, *α2δ-2* null animals exhibit severe cerebellar ataxia, spontaneous recurrent seizures, and a lowered seizure threshold (Snell, 1955; Ivanov et al., 2004; Danis et al., 2024). α2δ-2 is selectively expressed in many parvalbumin-expressing (PV+) inhibitory neurons throughout the brain, including cerebellar Purkinje cells and PV+ cells of the hippocampus (Barclay et al., 2001; Cole et al., 2005).

However, despite the important role that PV+ interneurons play in controlling hippocampal excitability (Trevelyan et al., 2006; Cammarota et al., 2013; Nahar et al., 2021), it is not known whether PV+ neuron-containing hippocampal microcircuits are affected by the loss of α2δ-2.

The hippocampal dentate gyrus is considered a gateway to seizure emergence, and dentate granule cells are highly activated during behavioral seizures in *α2δ-2* null mice (Krook-Magnuson et al., 2015; Danis et al., 2024). Thus, we used electrophysiology to assess how PV+ interneuron function in the dentate gyrus is altered in the absence of α2δ-2. Our results indicate that *α2δ-2* KO mice have substantially reduced PV+ interneuron-mediated inhibition, and that α2δ-2 is necessary for tight functional coupling of VGCCs to vesicle release in PV+ terminals.

## Methods

### Mice

Germline *α2δ-2* knockout mice (*Cacna2d2^tm1Svi^;* RRID:MGI:3055290) were generously provided by Drs. Sergey Ivanov and Lino Tessarollo at the National Cancer Institute (Ivanov et al., 2004). Mice homozygous for the targeted allele have the same null phenotype as the spontaneous *α2δ-2* null mutant mouse, *ducky^2J^* (Barclay et al., 2001). Mice were maintained as heterozygotes (no gross behavioral phenotype) and bred to create wildtype (WT) and knockout (KO) mice for experiments. Mice were genotyped using tissue samples at 1 week of age as previously reported (Beeson et al., 2020), and homozygous KO and WT littermates of both sexes were used for experiments at 3-6 weeks of age. Mice were housed in an AAALAC-approved facility on a 12hr light/dark cycle with constant access to food and water. Experiments were conducted in age-matched littermate controls whenever possible, and all experimental protocols involving mice were approved by the Portland Veterans Administration Institutional Animal Care and Use Committee.

For optogenetic stimulation, we used Cre-dependent Channelrhodopsin (ChR2) reporter mice obtained from Jax Laboratories (“ChR2”; B6.Cg-*Gt(ROSA)26Sor^tm32(CAG-COP4*H134R/EYFP)Hze^*/J; RRID: ISM_JAX:024109; Madisen et al., 2012). These mice were bred with heterozygous *α2δ-2* mice for two generations to obtain mice homozygous for ChR2 and heterozygous for *α2δ-2*. To easily visualize PV+ neurons for cell-attached recordings in acute slices, we crossed a PV interneuron-specific Cre line (“PV-Cre”; B6.129P2-*Pvalb^tm1(cre)Arbr^*/J; RRID: ISMR_JAX:017320; Hippenmeyer et al., 2005) with a Cre-dependent tdTomato fluorescent reporter mouse line (“tdT”; B6.Cg-*Gt(ROSA)26Sor^tm14(CAG-tdTomato)Hze^*/J; RRID: IMSR_JAX:007914, Madisen et al., 2010) and the *α2δ-2* mutant line, to obtain triple mutant PV Cre::tdT:: α2δ-2 mice. These mice were then crossed with our ChR2::α2δ-2 mice to produce offspring that were all heterozygous for each of these additional manipulations (PV-Cre^+/^::ChR2^+/-^::tdT^+/-^) and which included *α2δ-2* WT and *α2δ-2* KO littermates. These mice were used for optogenetic experiments at 3-6 weeks of age.

### Electrophysiology

Animals were terminally anesthetized via inhaled isoflurane followed by an intraperitoneal injection of 2% tribromoethanol (0.8 ml), and subsequently transcardially perfused with an ice-cold solution containing (in mM) 125 choline chloride, 7 MgCl2, 2.4 KCl, 1.25 NaH2PO42H2O, 0.5 CaCl2, 10 D-glucose, 1.3 Na-ascorbate, and 25 NaHCO3, 330 mOsm. Brains were sliced coronally into 300 μm sections in ice cold choline chloride solution on a Leica VT1200S vibratome and transferred to artificial cerebrospinal fluid (aCSF) containing (in mM) 125 NaCl, 25 NaHCO3, 1.25 NaH2PO4, 3 KCl, 25 D-glucose, 2 CaCl2, and 1 MgCl2, 310 mOsm at 35°C. After 30 minutes, slices were transferred to room temperature aCSF and recovered for an additional 30 minutes before experiments. All solutions were continuously bubbled with carbogen (95% O2 / 5% CO2).

Whole cell voltage clamp recordings were obtained using glass pipettes filled with a cesium-based internal solution containing (in mM) 113 CsGluc, 10 HEPES, 10 EGTA, 17.5 CsCl, 8 NaCl, 2 Mg-ATP, 0.3 Na-GTP, pH 7.28 - 7.24, 295-298 mOsm. Cell-attached recordings to measure optogenetic activation of PV-ChR+ interneurons used a potassium-based internal solution containing (in mM) 115 K Gluconate, 20 NaCl, 1.5 MgCl2, 5 HEPES, 10 K4BAPTA, 4 Mg-ATP, 0.3 Na-GTP, 10 phosphocreatine, 295 mOsm, pH = 7.28. Glass pipettes were pulled using a Sutter P-97 puller and tip resistances were between 3-5 MΩ for granule cell patching and 2-3 MΩ for PV+ interneuron patching. Slices were continuously superfused with carbogen-bubbled aCSF at 2 ml/min. Recordings were performed at room temperature (24°C) using an Axon Instruments MultiClamp 700B and acquired using custom-written software in IgorPro8 (WaveMetrics).

For granule cell recordings, cells in the outer half of the granule cell layer were chosen to avoid immature granule cells. All whole cell experiments were performed while cells were voltage-clamped to -70 mV. A 50 msec -10 mV voltage step was applied at the beginning of each sweep to calculate input resistance, series resistance, and cell capacitance. Cells were excluded if input resistance was >800 MΩ, if series resistance was > 20 MΩ or changed more than 15% from its initial value, or if holding current fell below -250 pA or changed more than 25% from its initial value. There were no differences in granule cell properties between genotypes in a subset of randomly chosen cells for direct comparison (input resistance: WT = 270 ± 22 MΩ, KO = 307 ± 23 MΩ, p = 0.242; series resistance WT = 16.5 ± 0.7 MΩ, KO = 16.7 ± 1.0 MΩ, p = 0.963, holding current WT = -107.0 ± 7.4 pA, KO = -90.2 ± 5.6 pA, p = 0.078, cell capacitance WT = 35.2 ± 1.9 pF, KO = 34.7 ± 1.7 pF, p = 0.837; WT n = 23 cells/10 animals, KO n = 22 cells/9 animals).

Electrically evoked synaptic responses were evoked every 10 seconds using an Iso-Flex stimulator (A.M.P.I.) and a bipolar electrode placed in the proximal inner molecular layer of the dentate gyrus, just above the granule cell layer. Stimulation intensity was adjusted to obtain an evoked response 60% of maximum (0.01 to 0.8 mA). When a stable baseline was reached, excitatory currents were pharmacologically isolated with application of 10 μM SR95531 (Abcam). At the end of each experiment, 10 μM NBQX (HelloBio) was applied to confirm that the remaining response was AMPA receptor-mediated; in all cases, no response remained. To calculate inhibitory responses, the averaged excitatory response obtained from 5 minutes of evoked recording was digitally subtracted from 5 minutes of the average baseline response using IgorPro8 to calculate the SR95531-sensitive (inhibitory) synaptic response. The maximum amplitudes from the excitatory and inhibitory averages were measured to determine the E/I ratio for each cell.

Optogenetically-evoked inhibitory post synaptic currents (oeIPSCs) from PV+ neurons in PV::ChR2 mice were obtained using 1 msec 470 nm light pulses delivered by a CoolLED pe300-ultra via a 60x water-immersion objective. Light intensity was measured using a thermal sensor power meter (Thorlabs) at increments between 0.38-7.97 mW/cm^2^. When a single light intensity was used, it is specified in the text/legend and all stimulation for that data set was conducted at the same intensity across slices and genotypes. To minimize direct depolarization of synaptic terminals by optogenetic stimulation, the visual field under the objective was moved ∼250 μm away from the patched cell along the granule cell layer. To determine paired pulse ratio, two LED pulses were delivered with a 100 msec inter-pulse interval, and the second oeIPSC amplitude was divided by the first oeIPSC amplitude. To estimate the readily releasable pool of vesicles, a 1.5 sec optogenetic stimulation train at 10 Hz was used. The cumulative responses were summed and graphed, and the last 5 values were used to fit a line and extrapolate it to the y-intercept, as previously described by Schneggenburger et al. (2002). Sets of responses for both paired pulses and stimulation trains were averaged from 20 trials after baseline stability was confirmed for 10 minutes and peak amplitudes were determined from these averages. For stimulation trains, peak amplitudes were graphed and used to determine the best fit for each cell using Microsoft Excel.

We used cell-attached recordings of PV+ interneurons to assess ChR2 function for both paired pulses and stimulation trains. After 5 minutes of a stable baseline, spiking responses were observed as narrow responses > -10 pA amplitude from baseline with spike waveforms. PV+ interneurons had a response rate of 100 ± 0% for all light intensities used in experiments in paired pulse and train stimulations for every trial (n = 3 cells/2 animals per genotype, p = 1.00).

To alter aCSF [Ca^2+^]ext to examine release probability, [Mg^2+^]ext was adjusted to maintain constant divalent ion concentration, specifically (in mM): 0.5 Ca^2+^/2.7 Mg^2+^ for low calcium, 2 Ca^2+^/1.2 Mg^2+^ for middle, and 3 Ca^2+^/0.2 Mg^2+^ for high calcium. Other aCSF concentrations were unchanged. To display inhibitory postsynaptic currents (IPSCs) for the calcium concentration experiments (Fig 8), traces from both genotypes were filtered for visualization using a 60 Hz low pass filter Hanning filter with smoothing window length of 30 from the IgorPro filter design and application function (WaveMetrics) to remove high frequency noise. Unfiltered traces were used for analysis, and filtering was strictly for better visualization and comparison of WT and KO traces in the figure.

For some experiments, the cell permeable calcium chelators EGTA-AM and BAPTA-AM (Cayman Chemical) were added to extracellular aCSF at 100 μM, similar to Hefft and Jonas (2005). EGTA-AM and BAPTA-AM were reconstituted in anhydrous DMSO (Fisher), and final DMSO concentration was limited to < 0.1%. aCSF was continuously recirculated for these experiments (1.2 ml/min with recirculation volume = 7 ml).

Spontaneous and miniature IPSCs (sIPSCs and mIPSCs) were recorded using 10 minutes of continuous acquisition in the presence of 10 μM NBQX (HelloBio). For miniature IPSCs, 500 nM TTX (Abcam) was added. sIPSCs and mIPSCs were quantified using custom-written scripts in IgorPro.

### Immunohistochemistry

For immunohistochemical processing (IHC), animals were terminally anesthetized using isoflurane and tribromoethanol (as above), transcardially perfused with 10 ml phosphate-buffered saline (PBS) followed by 10 ml 4% paraformaldehyde (PFA) in PBS (pH 7.3), and whole brains were post-fixed overnight. After fixation, brains were rinsed with PBS and sliced coronally on a Leica VT1000 vibratome into 100 μm sections. Slices were rinsed in PBS one more time before being blocked in a 10% goat serum solution containing PBS + 0.4% Triton X-100 (PBST) for one hour. Following blocking, slices were incubated in PBS containing 1.5% goat serum, 0.1% Triton-X100, and respective primary antibodies. For PV/VGAT colocalization, slices were incubated with primary antibodies against VGAT (rabbit, Synaptic Systems, 1:400, catalog #131-002; RRID: AB_887871) and PV (mouse, DHSB, 1:10, catalog # L114/3-5, RRID: AB_2877608) for 48 hours at 4°C. To assess tdT/ChR2 expression in PV cells, slices were incubated for 24 hours at 4°C with a primary antibody against PV (guinea pig, Synaptic Systems, 1:1000, catalog #195-308; RRID: AB_2927389) and an Alexa Fluor 488-conjugated antibody against GFP (rabbit, Invitrogen, 1:400, catalog #A-21311, RRID: AB_221477) to target ChR2-YFP. After 3x 5-minute washes in PBS, slices were incubated with respective secondary antibodies (Alexa Fluor 647 goat anti-rabbit, Jackson ImmunoResearch, 1:400, catalog #111-605-003, RRID: AB_2338072; Alexa Fluor 568 goat anti-mouse, Invitrogen, 1:400, catalog # A-11004, RRID: AB_2534072; Alexa Fluor 647 goat anti-guinea pig, Invitrogen, 1:400, catalog # A-21450, RRID: AB_2535867) for 24 hours at 4°C. tdT expression was assessed using endogenous fluorescence.

After 3x 5-minute PBS washes, slices were counterstained using 1:20,000 DAPI and then mounted on slides using Fluoromount-G (Southern Biotech). PV+ interneuron axons and terminals in the dentate granule cell layer were imaged using AIRYSCAN on a confocal laser-scanning microscope (LSM 980, Zeiss) with a 63x objective (Zeiss) at 2.5x zoom. Stacks of 13 images were taken at 0.17 μm Z-intervals. One image from each of two brain slices was captured for a total of 2 images per animal. Data from each animal was pooled and averaged.

VGAT/PV colocalization was determined using the Imaris imaging software surfaces function (ImarisColoc). Images were taken and file names were randomized and blinded before analysis. 3D surfaces were created of both the PV and VGAT channels. Minimum and maximum thresholds and filter values were set for each channel and applied across all images (PV = 750 min/1,500 max filtered at 500/1,200, VGAT 2,000 min/3,000 max, filtered at 500/2,000). VGAT colocalization with PV was determined using a distance of 0 μm between any VGAT and PV surface. The number of surfaces that fit this criterion was calculated using a custom RStudio program generously shared by Dr. Jennifer Jahncke. For density purposes, image volume was calculated from the imaging parameters; each image represented a tissue volume of 4410 μm^3^.

### Fluorescence in situ hybridization

*CACNA2D* isoform mRNAs were detected in fixed tissue using Hybridization Chain Reaction in situ mRNA staining (Molecular Instruments). *CACNA2D1* and *CACNA2D3* probes consisted of 20 probe sets targeting the full mRNA sequence of α2δ-1 and α2δ-3; *CACNA2D2* probes targeted the *CACNA2D2* exons (33-39) deleted in our *α2δ-2* KO mouse (Ivanov et al., 2004). Animals were fixed using transcardial perfusion of RNAse-free 4% PFA in PBS (pH 7.3) and whole brains were post-fixed for one hour. Brains were sliced coronally on a Leica vibratome to 100 μm thickness. Slices were mounted on SuperFrost Plus positively charged slides (FisherBrand) using RNAse free techniques and stained using slight modifications to the manufacturer’s protocol.

Briefly, slices were allowed to dry on the slides. To reduce autofluorescence, slides were then submerged in ice cold 1% NaBH4 (Sigma Aldrich) in PBS for 30 minutes, with fresh solution being applied every 10 minutes. Slides were subsequently rinsed in RNAse-free ice-cold PBS for 15 minutes, changing solution every 5 minutes, and then dehydrated in EtOH solutions as described in the Molecular Instruments HCR protocol.

Slides were prehybridized in hybridization buffer and then incubated in 0.4 pmol of each custom probe (parvalbumin (B5), *CACNA2D2* (B3), *CACNA2D1* (B4), and/or *CACNA2D3* (B2)) per 100 μL of hybridization buffer overnight in a heated, humidified chamber. The next day, SSCT washes were utilized as described in the HCR protocol and followed by application of probe amplification hairpins (488-B5, 567-B3 or -B2, 647-B4). After washing, TrueBlack lipofuscin binding solution (Biotium) was applied for 60 seconds to further reduce background, and slides were rinsed with EtOH and PBS before mounting in Fluoromount-G (Southern Biotech).

Images were taken on a spinning disc confocal microscope (Yokogawa CSU-W1) using a 60x objective (Zeiss) at a 2x zoom with Slidebook software (3i). Stacks of 10 images at Z intervals of 0.17 μm were obtained. Image files were blinded before analysis. Images cropped to 60 μm x 60 μm and an ROI was drawn around cells expressing PV+ RNA. Images of 10 different PV+ interneurons were taken across four dentate gyrus sections per animal. After identifying a PV+ interneuron using the PV RNA signal, each cell was manually scored for visible anti-*α2δ-1, -2*, or *-3* mRNA granules using the following criteria: a score of 0 when no RNA puncta were visible above background, 1 when RNA puncta were observed but the signal did not fill the cell area, 2 when RNA puncta clumped into groups, and 3 when the RNA signal was filling the cell body. After scoring, scores were averaged across each animal for comparison. Only *α2δ-2* WT animals scored higher than a 0 in the *α2δ-2* RNA expression levels, and neither genotype expressed higher than a 0 in *α2δ-1* or *α2δ-3*.

### Statistical Analysis

All statistical analysis was conducted using Prism (GraphPad) with significance set at p < 0.05. All bar graphs display mean ± SEM. Individual data points corresponding to the example traces shown in figures are highlighted in yellow on the corresponding graphs. To determine significance of comparisons between WT and KO groups, data normality was first assessed using a Shapiro-Wilks test. For normally distributed data, an unpaired Welch’s t-test was performed, otherwise a non-parametric Mann-Whitney test was performed. For light intensity curves and calcium chelator experiments, a two-way repeated measures ANOVA, or a mixed effects analysis in case of occasional missing values, was performed to compare WT and KO. For calcium concentration experiments, multiple unpaired t-tests were used. For *in situ* RNA characterization, a nested t-test was used to determine significance. For all experiments *p < 0.05, **p < 0.01, ***p < 0.001, ****p < 0.0001. For synaptic physiology, there were at least 5 animals per group in each set of experiments, and n is reported as number of cells. For all other experiments, n is reported as number of animals.

## Results

### Increased synaptic excitation to inhibition ratio in the dentate gyrus of α2δ-2 KO mice

The hippocampal dentate gyrus of *α2δ-2* KO mice demonstrates substantially increased granule cell activation in the absence of structural changes (Danis et al., 2024). As increased network excitability could result from reduced neuronal inhibition, we assayed inhibitory synaptic function in the dentate gyrus by assessing the ratio of synaptic excitation to inhibition (E/I ratio). We used electrical stimulation of the dentate inner molecular layer (IML), which contains both excitatory mossy cell inputs to granule cells, as well as inhibitory fibers that can be activated directly via electrical stimulation and indirectly via feed-forward mechanisms (Fig 1A). Evoked excitatory responses were isolated pharmacologically, and inhibitory responses were calculated via baseline subtraction (see Methods).

**Figure 1:**
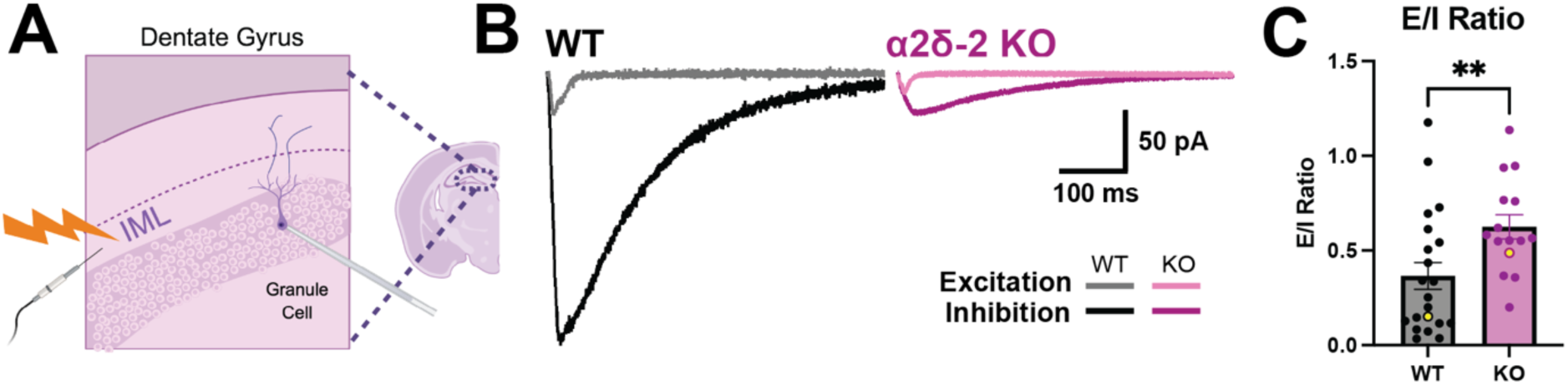
The ratio of synaptic excitation to inhibition (E/I) is increased in the dentate gyrus of *α2δ-2* KO animals. (A) Whole cell voltage clamp recordings of synaptic responses were obtained from dentate gyrus granule cells, in response to electrical stimulation of the inner molecular layer. (B) Example traces of isolated excitatory (light) and inhibitory (dark) evoked synaptic responses, obtained while keeping stimulation intensity constant for WT (left, black) and *α2δ-2* KO (right, magenta) slices. (C) The E/I ratio is increased in *α2δ-2* KO animals. Data from individual cells is represented by dots; yellow dots correspond to traces shown in B (WT = 0.37 ± 0.07, n = 21 cells; KO = 0.63 ± 0.06, n = 15 cells; p = 0.0096, Mann-Whitney test). Panel A created in BioRender. Ellingson, A.J. (2026) https://BioRender.com/3xkkpxu

The E/I ratio was significantly increased in *α2δ-2* KO mice (Fig 1B-D). This change could be due to increased excitatory mossy cell synaptic function, decreased inhibitory synaptic function, or decreased feedforward inhibition via a deficit in the mossy cell-mediated drive of inhibitory networks (Pinto et al., 2006). As hippocampal PV+ interneurons in CA1/3 selectively express α2δ-2, and PV+ neurons contribute to feedforward inhibition in the dentate gyrus (Cole et al., 2005; Willems et al., 2018), we pursued PV+ cells in the dentate gyrus for further investigation.

### PV+ interneuron-mediated synaptic inhibition is greatly diminished in α2δ-2 KO mice

To first confirm whether PV+ interneurons in the dentate gyrus also selectively express α2δ-2, we performed *in situ* hybridization for PV and the three brain-expressed α2δ isoforms in hippocampal sections. Consistent with prior observations in cortex and CA1/3 (Cole et al., 2005), α2δ-2 mRNA was clearly visualized in PV+ interneurons in WT mice (Fig 2A-B). We did not observe any α2δ-2 mRNA signal in non-PV+ cells, indicating that α2δ-2 is selectively expressed by PV+ interneurons in this region.

**Figure 2:**
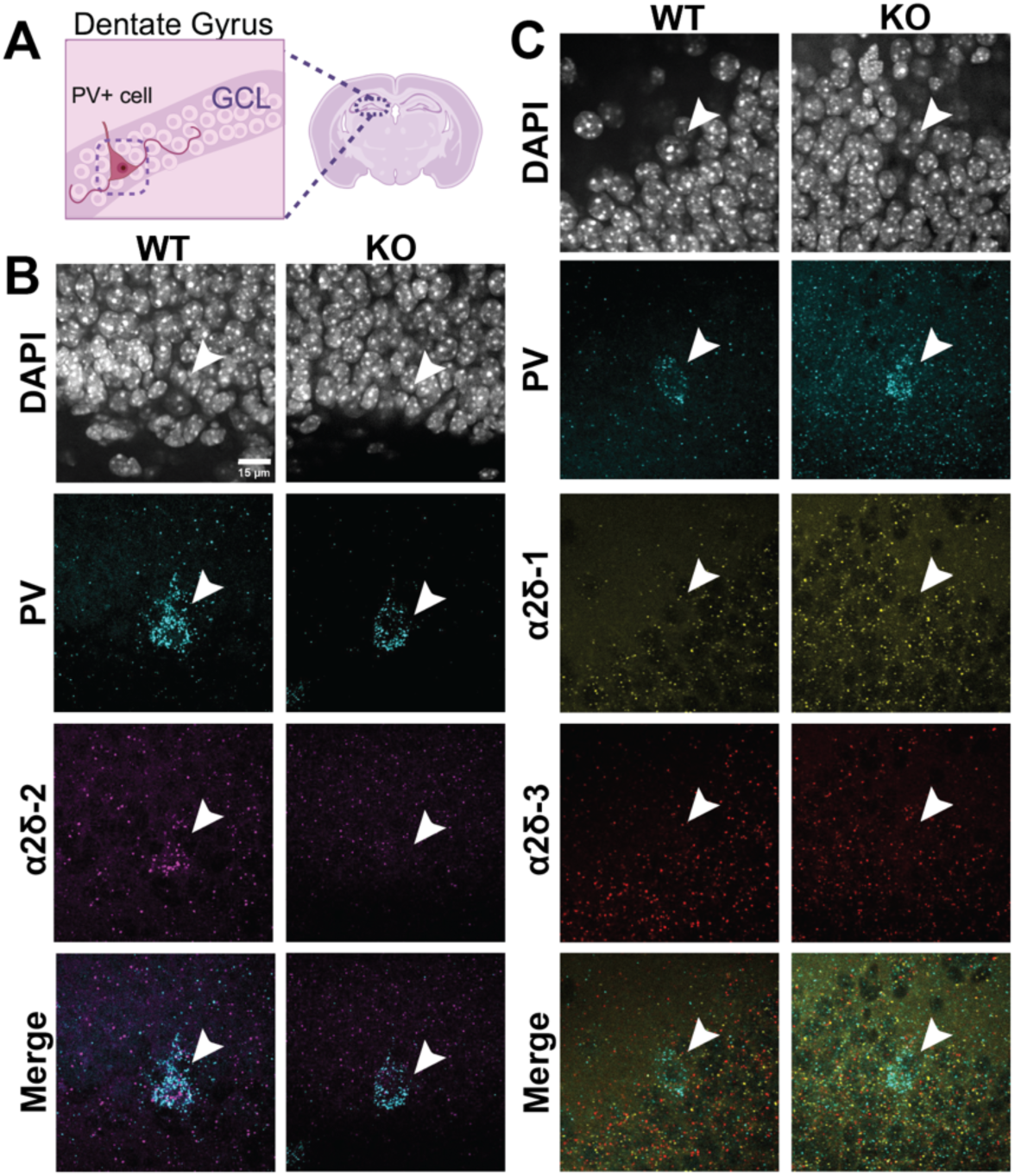
Dentate gyrus PV+ interneurons selectively express α2δ-2. (A) Fluorescent *in situ* mRNA hybridization and imaging was performed in the dentate granule cell layer, focusing on PV+ cell bodies. (B) α2δ-2 mRNA (magenta) was identified in PV+ interneurons (cyan, denoted by arrows) from WT but not *α2δ-2* KO animals (WT = 1.2 ± 0.3 score for presence of RNA, n = 3; KO = 0 ± 0, n = 3; p = 0.026, nested t-test); α2δ-2 granules were not detected in non-PV+ cells. Granule cell layer nuclei were visualized with DAPI (white). Scale bar = 15μm. (C) α2δ-1 and α2δ-3 mRNA (yellow and red, respectively) was not detected in PV+ interneurons in either WT or KO animals (WT = 0 ± 0, n = 3; KO = 0 ± 0, n = 3, p = 1.0 for both α2δ-1 (yellow) and α2δ-3 (red)), same scale. Panel A created in BioRender. Ellingson, A.J. (2026) https://BioRender.com/huvfthc

Conversely, although mRNA granules for α2δ-1 and -3 were present in other cell types in the dentate gyrus (including putative dentate granule cells and hippocampal mossy cells, based on location), these isoforms were not detected in PV+ cells. In *α2δ-2* KO mice, α2δ-2 mRNA signal was absent from PV+ cells, consistent with gene deletion, but mRNA for other α2δ isoforms was still not detected (Fig 2C). Thus, α2δ-2 is selectively expressed by PV+ cells in the dentate gyrus, and loss of α2δ-2 does not result in clear upregulation of other α2δ isoforms.

As there is no difference in the PV+ interneuron density in the dentate gyrus between WT and *α2δ-2* KO mice (Danis et al., 2024), loss of PV+ neurons in KO mice could not explain the increased E/I ratio. Thus, we assessed PV+ interneuron function using genetically modified mice expressing channelrhodopsin (ChR2) in PV+ interneurons. Driver and reporter mice were crossed with *α2δ-2* mutant mice to generate littermate PV-Cre::ChR2::tdT::α2δ-2 WT and KO mice (see Methods), to facilitate visual identification and optogenetic activation of PV+ neurons in both *α2δ-2* genotypes.

We first assessed Cre-dependent reporter expression in WT and *α2δ-2* KO PV-Cre::ChR2::tdT::α2δ-2 mice to confirm effective labeling of PV+ cells. In both WT and KO mice, we observed selective and specific expression of both reporters in PV+ cells, with high penetrance and little off-target reporter expression (Fig 3A). Importantly, we found no difference in the density of reporter-expressing PV+ cells between WT and KO mice (Fig 3B), as expected from prior observations (Danis et al., 2024). As cerebellar Purkinje cells lacking α2δ-2 have altered spiking dynamics (Walter et al., 2006; Beeson et al., 2022), we also wanted to confirm that we would be able to effectively drive PV-mediated inhibition using optogenetics in *α2δ-2* KO mice. Using cell attached recordings from PV-ChR2+ interneurons, PV+ cells from both genotypes reliably fired action potentials in response to paired pulse and train activation patterns at all light intensities (Fig 3C-D). PV+ interneurons often fire sets of two or more action potentials in response to synaptic depolarization (Milicevic et al., 2024) and PV+ interneurons from WT and KO mice also fired ‘doublets’ in response to optogenetic stimulation, at similar rates between genotypes (Fig 3C-D).

**Figure 3:**
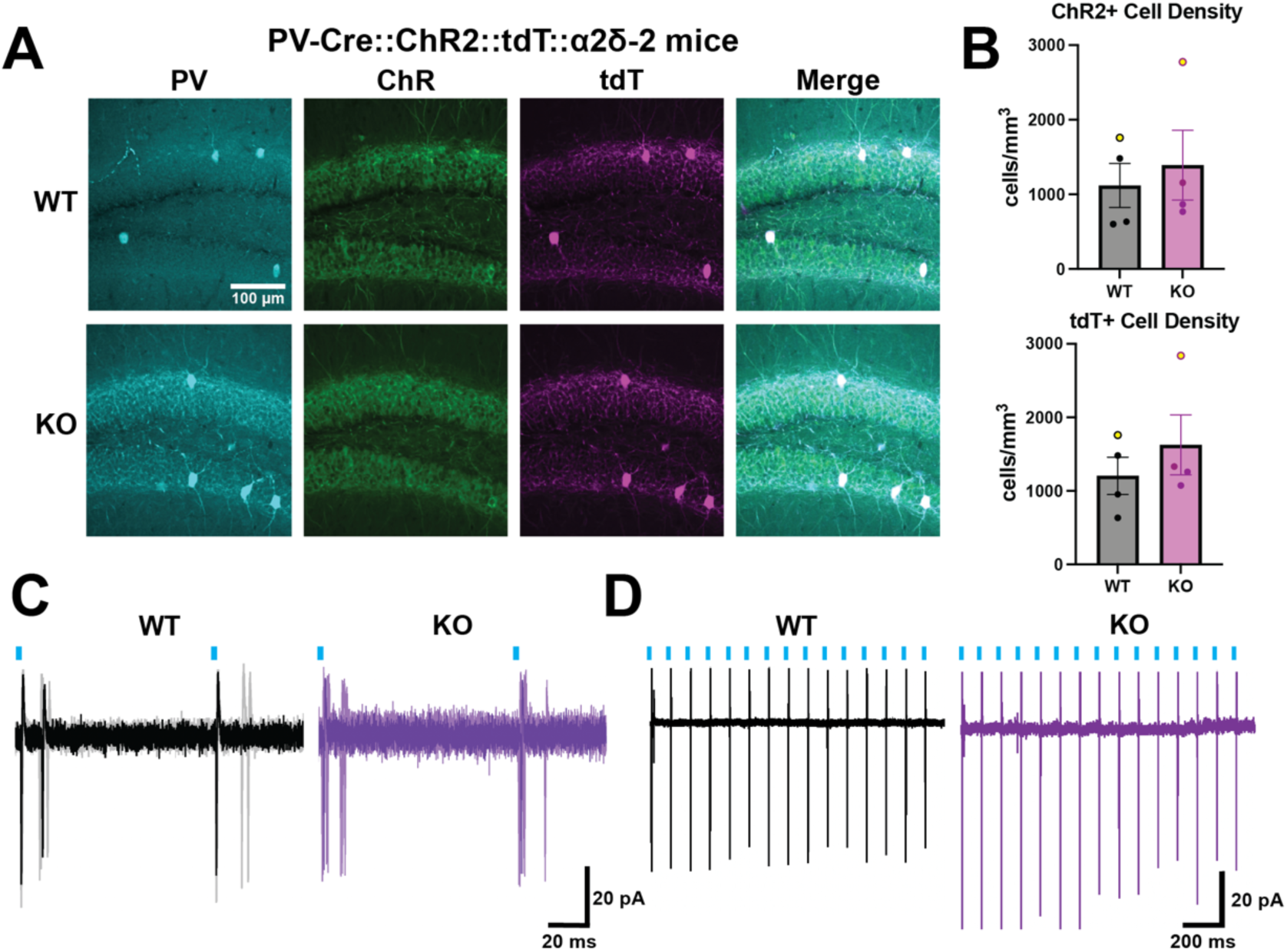
Channelrhodopsin reporter expression and function in PV+ interneurons from WT and *α2δ-2* KO animals. (A) Hippocampal slices from PV-Cre::ChR2::tdT::α2δ-2 animals were stained for PV and ChR2-YFP, with endogenous tdTomato fluorescence used to determine cell-specific expression. Images from representative WT (top) and *α2δ-2* KO (bottom) sections shown. (B) PV-ChR2+ and PV-tdT+ neuron densities were not different between genotypes (n = 4 WT, n = 4 KO mice; PV-ChR2+ p = 0.64, PV-tdT+ p = 0.42). Yellow dots correspond to mice shown in A. (C, D) Cell attached recordings from PV+ interneurons during optogenetic paired pulse (C) and 10 Hz train (D) stimulation. No firing failures were observed in either genotype, and both genotypes had similar doublet rates across increasing light intensities (0 ± 0% doublets for both WT and KO at intensities between 0.38 and 0.75mW/cm^2^; WT = 23.3 ± 23.3%; KO = 20.1 ± 6.7% at 1.35mW/cm^2^; and 100 ± 0% doublets at light intensities between 2.45 and 7.97; n = 3 cells/2 animals per genotype).

To assess presynaptic function of dentate gyrus PV+ interneurons, we recorded from postsynaptically connected granule cells while optogenetically driving PV+ neurons in acutely prepared hippocampal slices from PV-ChR2 *α2δ-2* WT and KO mice (Fig 4A). Optogenetically-evoked inhibitory post synaptic currents from PV+ interneurons (PV-oeIPSCs) were 72% smaller in granule cells from *α2δ-2* KO mice than in those from WT littermates (Fig 4B-C). Increased light intensity evoked larger PV-oeIPSCs in both genotypes, but responses in KO mice remained smaller than WT littermates across all light intensities (Fig 4D). PV-oeIPSC decay kinetics were 20% faster in *α2δ-2* KO animals (Fig 4E) (WT tau = 20.6 ± 1.5 msec, n = 28; KO tau = 16.1 ± 1.2 msec, n = 26; p = 0.02, Welch’s unpaired t-test). Reduced PV-oeIPSCs were not associated with an overall loss of inhibitory synaptic innervation, as assessed by miniature or spontaneous IPSCs (Fig 5) and thus appeared to result from reduced functional PV+ interneuron-granule cell synapses in *α2δ-2* KO mice.

**Figure 4:**
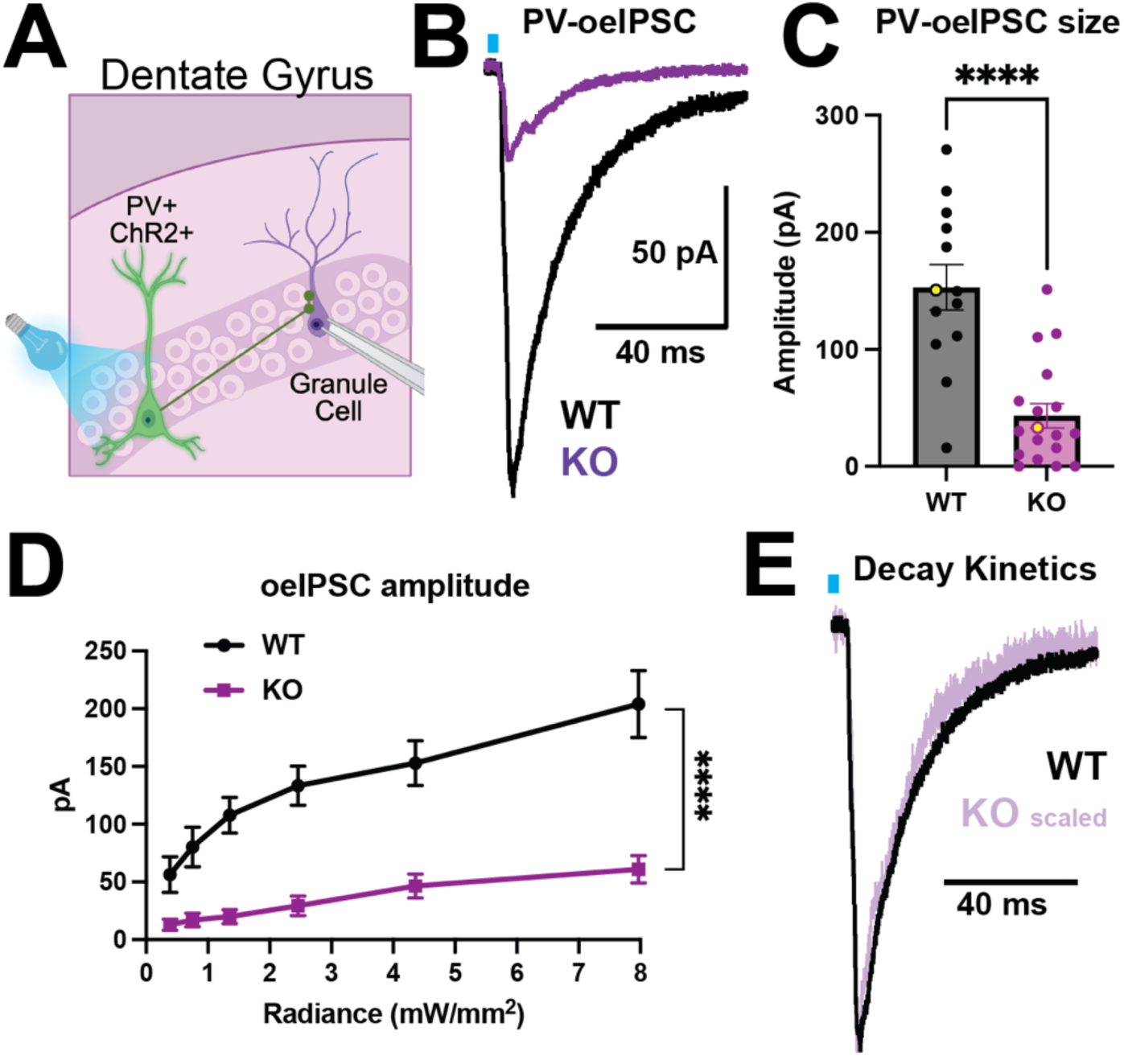
PV+ interneuron to granule cell synaptic function is dramatically reduced in *α2δ-2* KO animals. (A) Experimental schematic. PV+ cells were optogenetically activated and synaptic responses were recorded from dentate granule cells via whole cell voltage clamp electrophysiology. (B) Example optogenetically-evoked PV IPSCs (PV-oeIPSCs) recorded from WT and *α2δ-2* KO granule cells at a light intensity of 4.4 mW/mm^2^. (C) PV-oeIPSC amplitudes are reduced in *α2δ-2* KO granule cells. (WT = 153 ± 19.3pA, n = 13; KO = 43.4 ± 10.3pA, n = 18; p < 0.0001, unpaired t-test), yellow dots correspond to traces shown in B. (D) Reduced PV-oeIPSC amplitude in *α2δ-2* KO mice is independent of stimulation light intensity (n = 18, p < 0.0001 for all conditions, two-way ANOVA). (E) α2δ-2 KO PV-oeIPSC decay is slightly but significantly faster. Panel A created in BioRender. Ellingson, A.J. (2026) https://BioRender.com/mp0whdo

**Figure 5:**
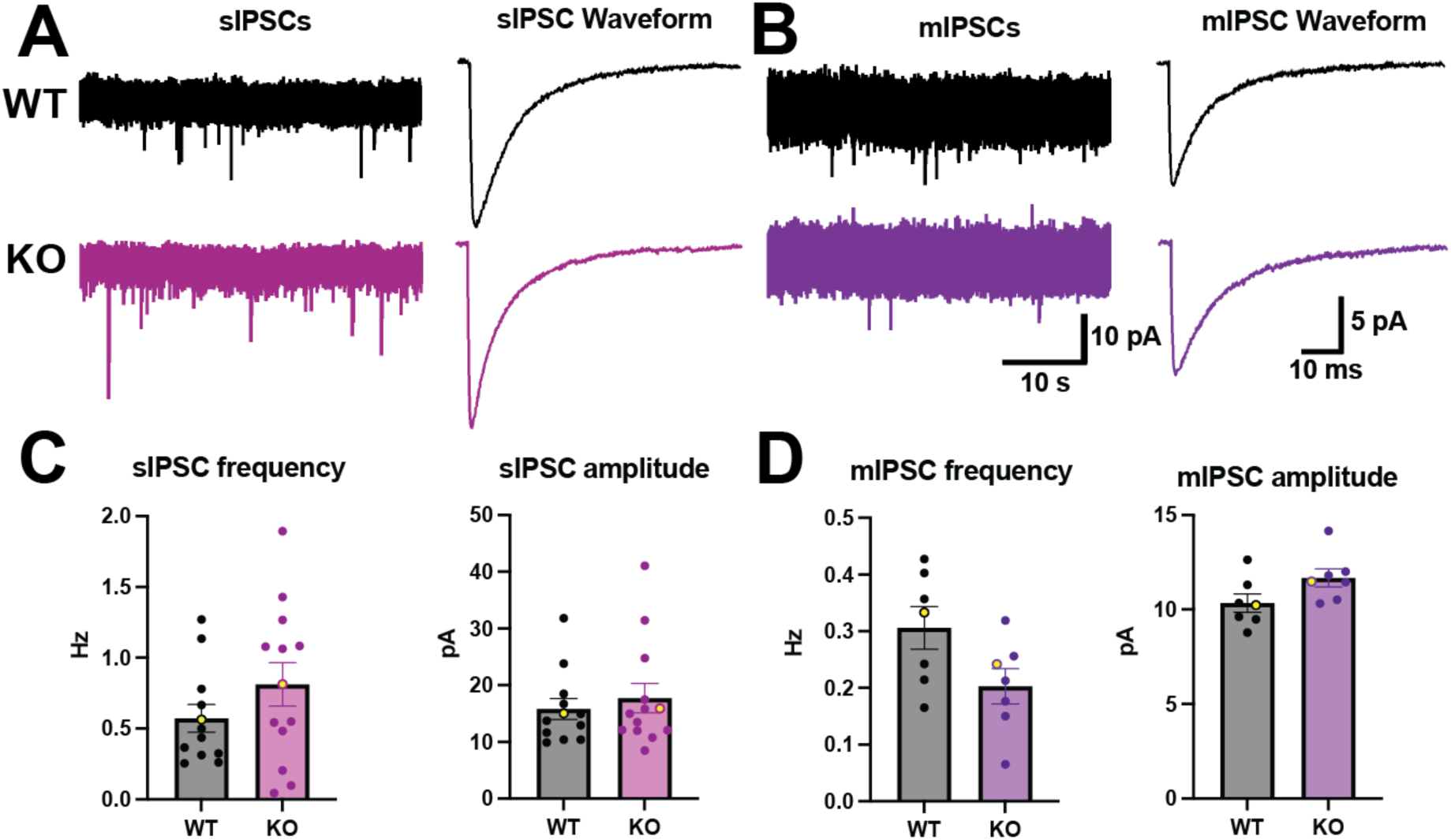
Spontaneous and miniature inhibitory postsynaptic currents are unchanged in α2δ-2 KO granule cells. (A, B) Example recordings of spontaneous (A) and miniature (B) IPSCs from WT and *α2δ-2* KO granule cells. (C) sIPSC frequency and amplitude are unchanged in *α2δ-2* KOs (Frequency: WT = 0.57 ± 0.10 Hz, n = 12; KO = 0.81 ± 0.15 Hz; n = 13; p= 0.2; Amplitude: WT = 15.8 ± 1.9 pA, n = 12; KO = 7.7 ± 2.6 pA, n = 13; p = 0.56; Welch’s t-tests). (D) mIPSC frequency and amplitude are unchanged in *α2δ-2* KOs (Frequency: WT = 0.31 ± 0.04 Hz, n = 7; KO = 0.20 ± 0.03 Hz; n = 7; p = 0.07; Amplitude: WT = 10.3 ± 0.5 pA, n = 7; KO = 11.7 ± 0.5, n = 7; p = 0.06; Welch’s t-tests). Yellow dots for each graph correspond to traces shown in A and B.

### α2δ-2 KO mice have fewer PV+ synaptic terminals in the dentate granule cell layer

α2δ proteins modulate the assembly and composition of synapses (Eroglu et al., 2009; Geisler et al., 2019; Schöpf et al., 2021). To assess whether a reduced number of PV+ inhibitory synapses could explain the decreased PV-oeIPSCs in *α2δ-2* KO mice, we performed immunohistochemistry on fixed sections of the dentate gyrus. We defined putative PV+ axon terminals as punctate structures where PV colocalized with the inhibitory synapse marker VGAT (Fig 6; see Methods).

**Figure 6:**
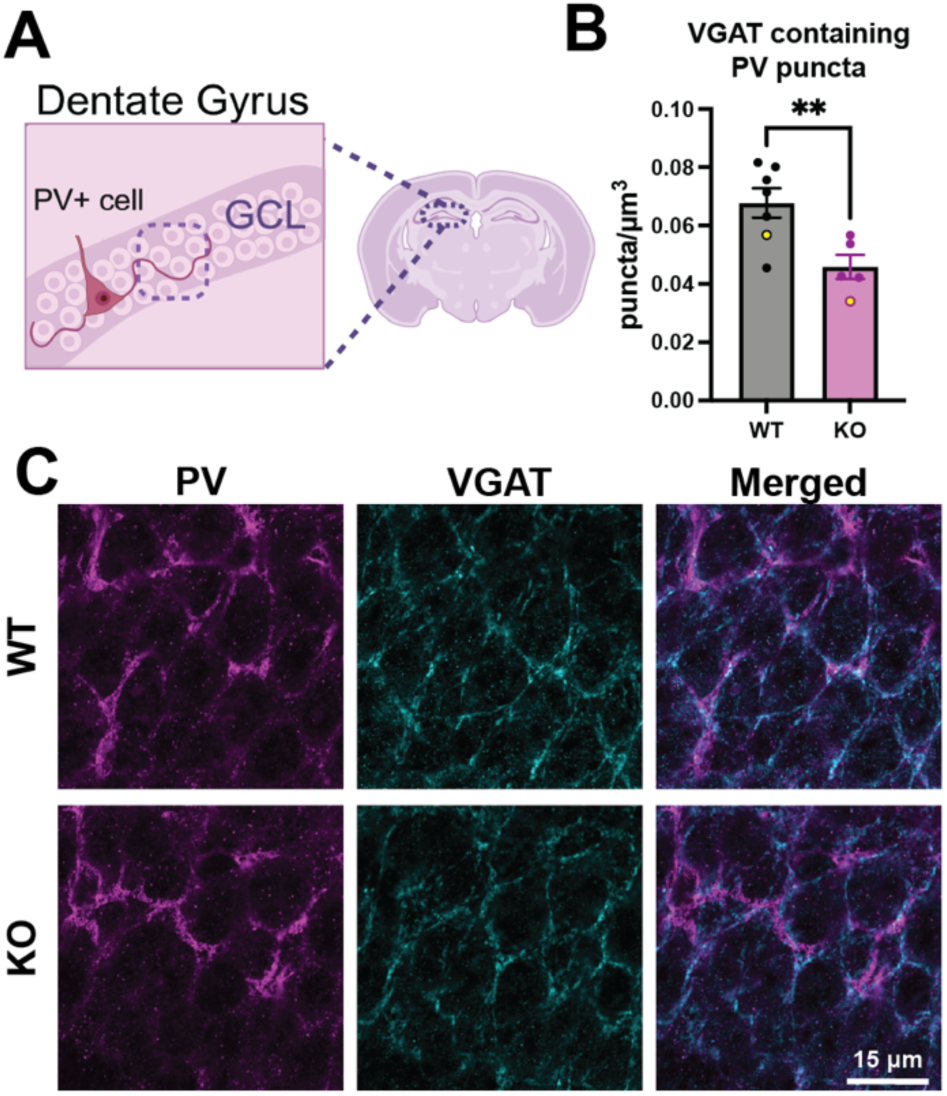
PV+ synaptic terminal density is reduced in *α2δ-2* KO dentate granule cell layer. (A) Schematic demonstrating region imaged for analysis. (B) *α2δ-2* KO mice have reduced density of VGAT puncta containing PV (WT = 0.068 ± 0.005 puncta/μm^3^, n = 7 mice; KO = 0.046 ± 0.004 puncta/μm^3^, n = 5 mice; p = 0.007, Welch’s t-test). Yellow dots correspond to the examples shown in C. (C) Example dentate granule cell layer images stained with antibodies to PV (magenta) and VGAT (cyan) from WT and *α2δ-2* KO mice. Panel A created in BioRender. Ellingson, A.J. (2026) https://BioRender.com/vykbbap

There was a 32% reduction in the density of VGAT puncta that colocalized with PV+ axons in *α2δ-2* KO dentate granule cell layer (Fig 6A-C). There was also a reduction in the volume of PV+ axon/terminal labeling in KO mice as a percentage of total image volume (WT = 0.093 ± 0.012%, n = 7; KO = 0.058 ± 0.007%, n = 5, p = 0.037). There was no change in the VGAT puncta density (WT = 0.245 ± 0.027 puncta/μm^3^; KO = 0.218 ± 0.013 puncta/μm^3^, p=0.406) or average volume of each VGAT punctum (WT = 0.53 ± 0.07 μm^3^; KO = 0.63 ± 0.25 μm^3^; p = 0.729). Therefore, although the overall density of inhibitory synapses (defined immunohistochemically) was unchanged in KO mice, there appeared to be a reduction in PV+VGAT+ synaptic terminals.

Although the density of PV+ neurons in the dentate gyrus was unchanged, a decrease in the density of PV+ inhibitory synapses could contribute to the reduced PV-oeIPSC amplitude in *α2δ-2* KO mice. However, the ∼30% reduction in PV+ inputs does not account for the ∼70% reduction in PV-oeIPSC size (Fig 4), suggesting additional functional differences.

### Probability of vesicle release and readily releasable vesicle pool size are reduced in PV+ interneurons in α2δ-2 KO mice

α2δ-1 couples presynaptic calcium entry to vesicle release in cultured pyramidal cells, and altered presynaptic α2δ expression can alter the probability of vesicle release (Hoppa et al., 2012). To assess a potential presynaptic role of α2δ-2 in inhibitory PV+ interneurons, we assayed the probability of vesicle release from PV+ cells using optogenetic paired pulse stimulation of PV+ interneuron axons (Figure 7A-B). The paired pulse ratio (PPR) was increased in *α2δ-2* KO mice, consistent with a reduction in probability of GABA release from PV+ interneurons lacking α2δ-2 (Figure 7C).

**Figure 7:**
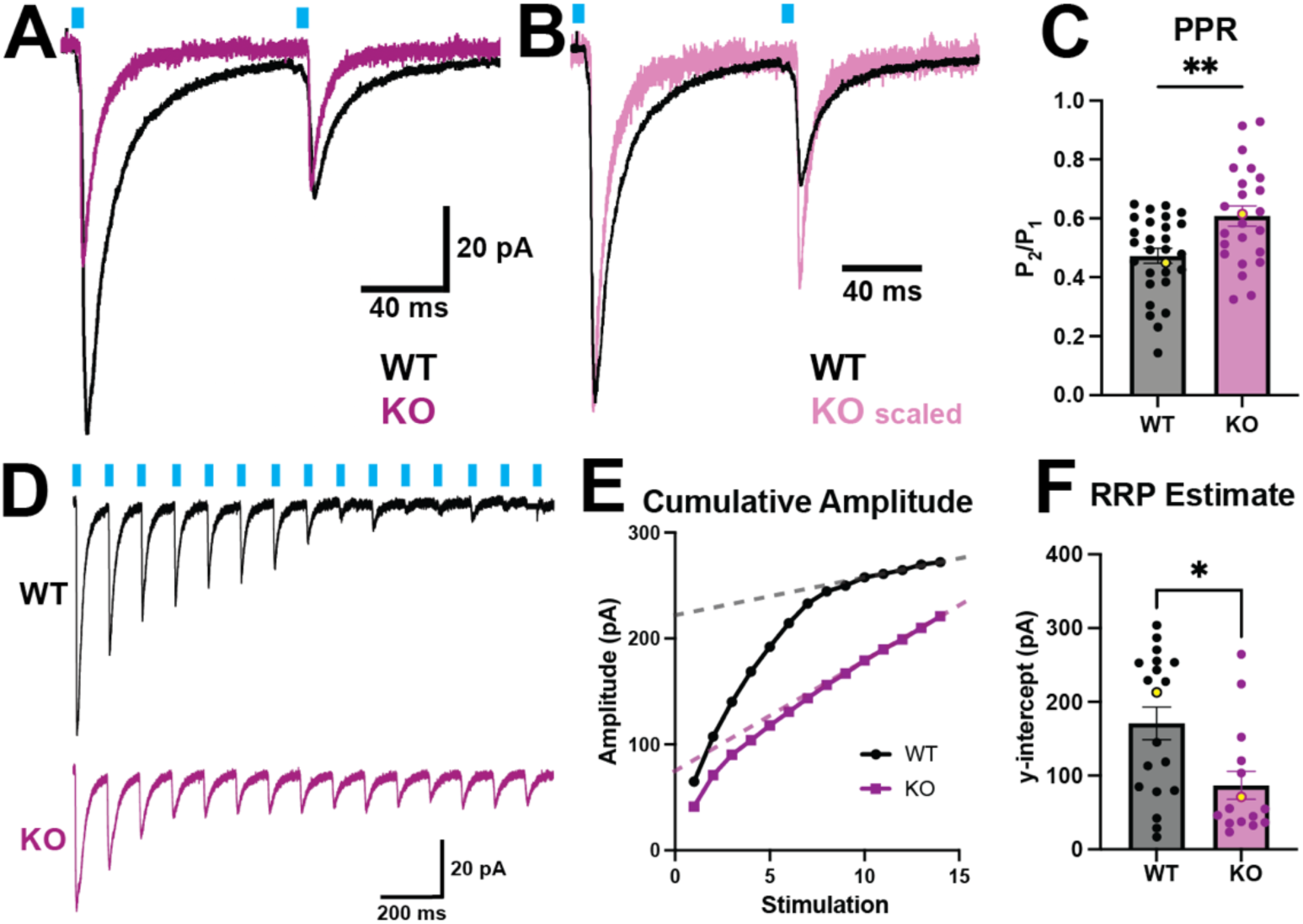
*α2δ-2* KO PV+ interneurons have a reduced probability of vesicle release and readily releasable vesicle pool size. (A) Example PV-oeIPSCs from WT (black) and *α2δ-2* KO (magenta) granule cells recorded during paired pulse optogenetic stimulation, overlaid for comparison. (B) The same traces from (A), scaled to align the peaks of the first response. (C) *α2δ-2* KO animals have a higher paired pulse ratio (PPR; WT = 0.47 ± 0.03, n = 28; KO = 0.61 ± 0.03, n = 24; p = 0.0027, Welch’s t-test). Yellow dots correspond to ratios from traces shown in A. (D) PV-oeIPSCs from WT (black) and *α2δ-2* KO (magenta) granule cells recorded during optogenetic stimulation trains at 10 Hz. (E) Cumulative amplitude plot for the example traces shown in D; the last 5 points of the plot were used to determine a line of best fit and its y-intercept. (F) Readily releasable pool (RRP) size was smaller in *α2δ-2* KO granule cells (WT = 171 ± 22 pA, n = 19; KO = 87 ± 19 pA, n = 15; p = 0.015, Mann-Whitney test). Yellow dots correspond to the examples shown in D and E.

The size of the readily releasable pool of vesicles (RRP) contributes to the probability of presynaptic vesicle release in the cerebellum (Vaden et al., 2019). To assess the RRP in dentate gyrus PV+ interneurons, we used a train of 15 optogenetic stimuli delivered at 10 Hz (Figure 7D). Using the method described by Schneggenburger et al. (2002) (see Methods), we estimated the RRP size independent of the vesicle recycling and release rate by extrapolating a line corresponding to the steady-state increase rate of cumulative PV-oeIPSC amplitude back to the y-intercept (Fig 7E). With this approach, the RRP in *α2δ-2* KO mice was reduced by 49% (Fig 7F). Together, the reduction in release probability and RRP size suggest that reduced GABA release from PV+ interneuron terminals lacking α2δ-2 contributes to the reduced PV-oeIPSC.

### Coupling of presynaptic Ca^2+^ entry to GABA release from PV+ interneurons is diminished in α2δ-2 KO mice

At excitatory synapses, α2δ proteins traffic VGCCs to the cell surface and presynaptic sites, and couple VGCC-mediated Ca^2+^ influx to vesicular release (Davies et al., 2010; Hoppa et al., 2012; Schöpf et al., 2021). To assess whether decreased VGCC abundance at PV+ interneuron terminals could account for reduced PV+ interneuron GABA release in KO mice, we altered Ca^2+^ influx into presynaptic terminals by systematically varying extracellular calcium concentrations while maintaining a constant extracellular divalent ion concentration (see Methods). We optogenetically evoked PV-oeIPSCs at a constant light intensity across slices while recording from dentate gyrus granule cells (Fig 8A). Although PV-oeIPSC amplitudes in each genotype increased with calcium concentrations, they did not converge (Fig 8B). Higher extracellular calcium concentrations did cause convergence of PPR between genotypes (Fig 8C-D), but the persistently smaller PV-oeIPSCs suggest that reduced VGCCs in PV+ terminals may not sufficiently explain the reduced PV-oeIPSCs in KO mice.

**Figure 8:**
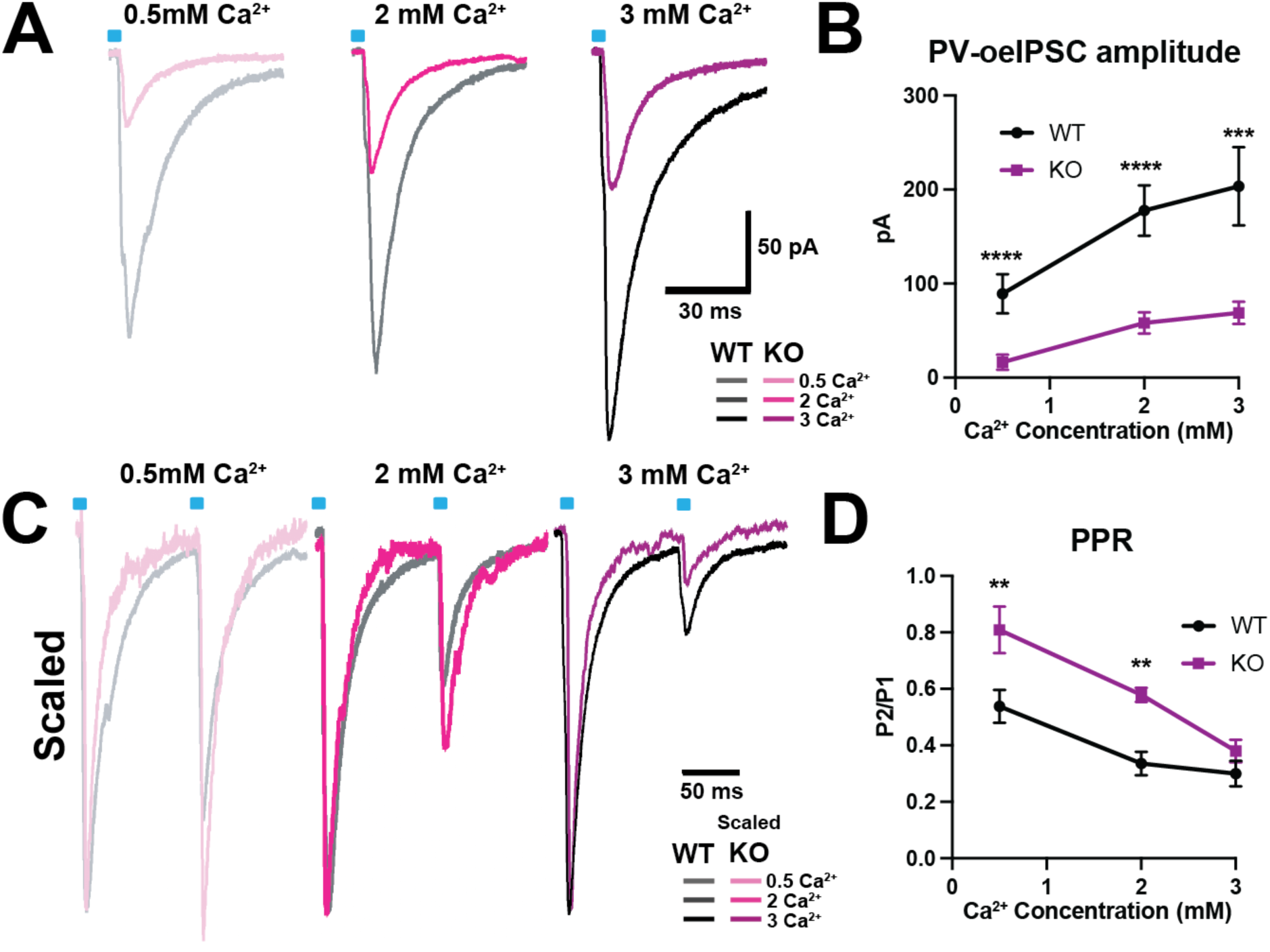
Increased extracellular calcium does not rescue PV-oeIPSC amplitudes in *α2δ-2* KO granule cells. (A) Example PV-oeIPSC averages recorded from WT (grey) and *α2δ-2* KO (magenta) granule cells in different extracellular calcium concentrations. (B) Increased extracellular calcium does not rescue PV-oeIPSC amplitude in *α2δ-2* KO granule cells (p < 0.001 for **** and p < 0.005 for *** for multiple unpaired t-tests). (C) PV-oeIPSCs from WT (grey) and *α2δ-2* KO (magenta) granule cells during paired pulse optogenetic stimulation in different extracellular calcium concentrations, overlaid and scaled to the first oeIPSC amplitude for each concentration. (D) PPR is restored in *α2δ-2* KO granule cells in higher extracellular calcium concentrations (n = 13; p = 0.0005 for 2-way ANOVA; p < 0.001 for **** and p < 0.005 for *** for multiple unpaired t-tests).

GABA release from PV+ interneurons in the hippocampus is tightly coupled to VGCC-mediated calcium entry, as evidenced by its insensitivity to the slow calcium chelator EGTA-AM and weak sensitivity to the fast calcium chelator BATPA-AM (Hefft and Jonas, 2005). If α2δ-2 functionally couples Ca^2+^ entry to vesicle release in PV+ cells, reduced Ca^2+^-release coupling could explain the reduced release probability in *α2δ-2* KO PV+ interneuron terminals.

To examine this, we optogenetically stimulated PV+ interneurons while recording from postsynaptic granule cells, and then bath applied the membrane-permeable calcium chelators EGTA-AM or BAPTA-AM (Fig 9). Although PV-oeIPSCs in WT mice were insensitive to EGTA-AM, PV-oeIPSCs in *α2δ-2* KO animals were reduced by EGTA-AM (Fig 9A-B). Furthermore, while BAPTA-AM did cause a small reduction in PV-oeIPSC amplitude in WT mice, this inhibition was greater in recordings from *α2δ-2* KO mouse slices (Fig 9C-D). Together, these data demonstrate increased sensitivity of GABA release from PV+ terminals to Ca^2+^ chelation in *α2δ-2* KO mice, indicating that the tight nanodomain coupling of VGCCs to release in PV+ terminals was diminished.

**Figure 9:**
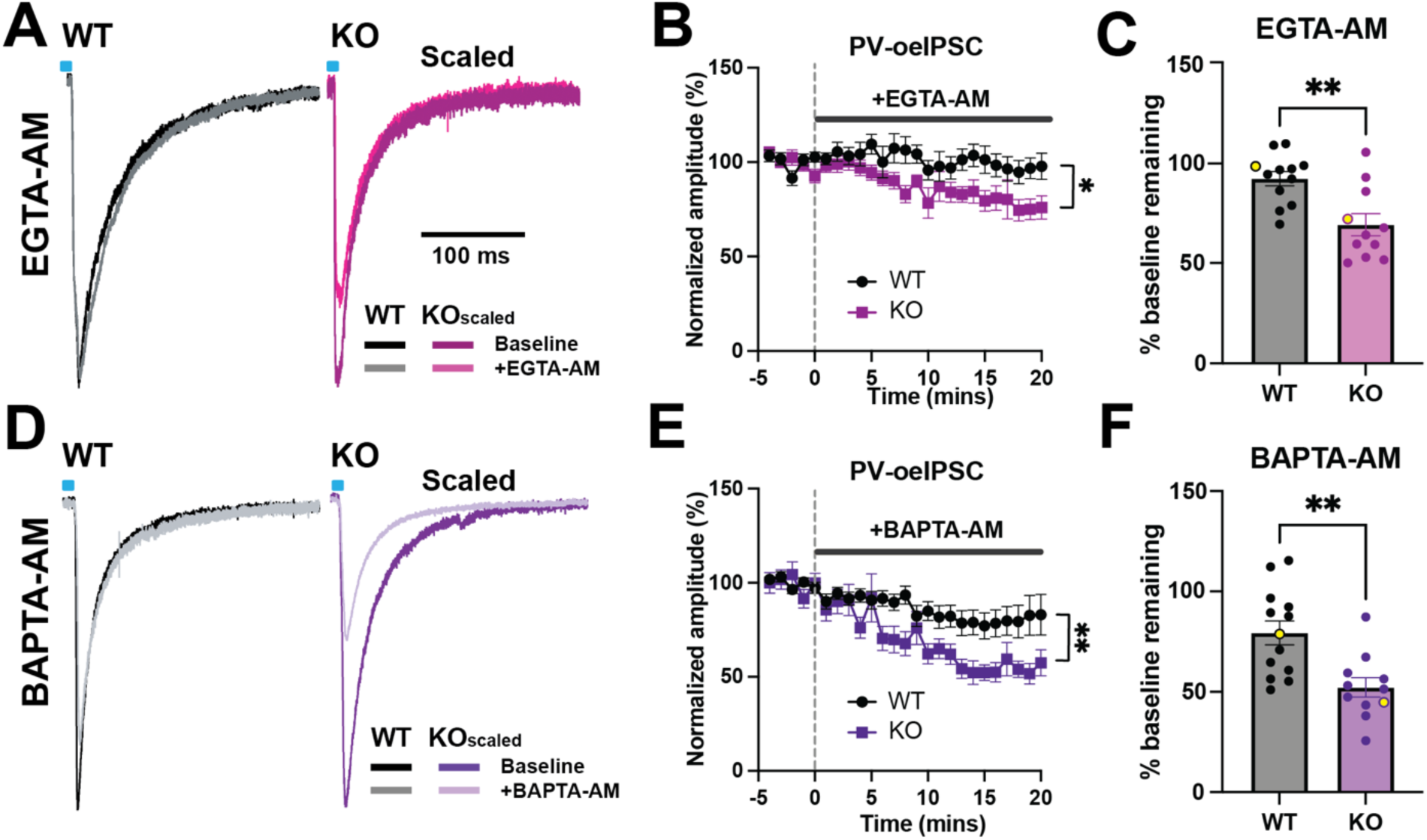
PV+ synaptic transmission is less tightly coupled to calcium entry in *α2δ-2* KO animals. (A) Example PV-oeIPSCs from WT and *α2δ-2* KO granule cells before and after application of the cell-permeable calcium chelator EGTA-AM; PV-oeIPSCs were for each cell were scaled to make amplitudes of the baseline responses equivalent. (B) PV-oeIPSC amplitudes from WT and *α2δ-2* KO granule cells during bath application and cell uptake of EGTA-AM, beginning at time = 0 (n = 12 WT, n = 11 KO; p = 0.02, mixed effects analysis). (C) PV-oeIPSCs are sensitive to EGTA-AM in *α2δ-2* KO granule cells, but not granule cells from WT mice (residual amplitudes relative to baseline: WT = 92.0 ± 3.6%, n = 10; KO = 69.1 ± 5.5%, n = 7; p = 0.003, Welch’s t-test). Yellow dots correspond to traces in A and their diary plots in B. (D) Peak-scaled PV-oeIPSCs before and after BAPTA-AM. (E) PV-oeIPSC amplitudes from WT and *α2δ-2* KO granule cells during bath application of BAPTA-AM (n = 13 WT, n = 11 KO; p = 0.005, mixed effects analysis). (F) *α2δ-2* KO PV-oeIPSCs are more sensitive to BAPTA-AM (residual amplitudes: WT = 79.4 ± 6.0%, n = 13; KO = 52.2 ± 0.9%, n = 11; p = 0.002, Welch’s t-test). Yellow dots correspond to traces shown in D and their diary plots show in E.

Thus, we conclude that α2δ-2 contributes to functional coupling between presynaptic Ca^2+^ entry and GABA release from PV+ interneurons.

## Discussion

Selective expression of a single α2δ isoform, α2δ-2, in PV+ interneurons makes them particularly susceptible to mutations in the *α2δ-2* gene. This presents the unique opportunity to use *α2δ-2* mutant mice to illuminate the functional roles of α2δ proteins in PV+ neurons. Together with electrophysiological studies of PV+ Purkinje cells in the cerebellum (Barclay et al., 2001; Walter et al., 2006; Beeson et al., 2020, 2022), these mice have extended our understanding of how α2δ proteins control VGCC-mediated signal transduction. Additionally, they also provide insights into how signaling cascades in subpopulations of brain neurons contribute to neurological phenotypes.

### α2δ proteins functionally couple VGCCs to downstream effectors

Our findings support a model in which α2δ proteins critically couple calcium entry to downstream effector mechanisms (Hoppa et al., 2012; Beeson et al., 2022). Although α2δs are important for functional coupling, α2δs also mediate trafficking of VGCCs to the cell surface and to the presynaptic terminal (Shistik et al., 1995; Hoppa et al., 2012; Milanick et al., 2025), which could also contribute to the functional deficits in *α2δ-2* mutant mice and explain the normalization of paired pulse ratio with an increase in [Ca^2+^]ext. It is not clear whether the contribution of α2δs to presynaptic terminal trafficking and nanodomain coupling is independent of their role in cell surface trafficking, as surface trafficking and subsequent (sub)synaptic localization are separately regulated for other synaptic proteins, such as presynaptic neurexins and postsynaptic AMPA receptors (Chen et al., 2000; Fairless et al., 2008; Neupert et al., 2015). Although our data cannot explicitly confirm whether *α2δ-2* KO mice have fewer VGCCs at PV+ presynaptic terminals, that alone could not fully explain the increased sensitivity of *α2δ-2* KO PV-oeIPSCs to EGTA-AM. Specifically, if there were fewer VGCCs in PV+ terminals, but the VGCCs present were otherwise appropriately localized relative to calcium-dependent release machinery, tight functional coupling (and EGTA-AM resistance) would be preserved. Thus, α2δ proteins could play roles in three separate subcellular events, namely: surface trafficking, synaptic localization, and subsequent nanodomain coupling.

As α2δs are exclusively extracellular (Jay et al., 1991), how might they mediate coupling between VGCCs and intracellular effector mechanisms? α2δ proteins interact with several extracellular binding partners via a metal ion-dependent adhesion site (MIDAS) domain on the α2 subunit (Cantí et al., 2005). In hippocampal cultures, α2δ constructs with mutated MIDAS domains rescue VGCC trafficking to the cell surface, but are unable to rescue VGCC-release coupling (Hoppa et al., 2012). This distinction suggests that extracellular interactions may be critical to coupling and also separable from surface trafficking functions. As the α2δ MIDAS domain interacts with synaptic adhesion proteins (Whittaker and Hynes, 2002; Lacy et al., 2004; Springer, 2006), its role in facilitating coupling might rely more on proper subcellular, or even subsynaptic, localization of the VGCCs in relation to intracellular effector mechanisms, rather than by directly bridging VGCC-effector interactions. α2δ interactions might involve binding to presynaptic neurexin molecules (Brockhaus et al., 2018), which may serve to localize VGCCs to specific terminal domains. Alternatively, α2δ proteins engage in trans-synaptic interactions, which might serve to align VGCCs with specific pre- and post-synaptic elements (Fell et al., 2016; Geisler et al., 2019). Thus, extracellular interactions could properly localize VGCCs to vesicular release sites at PV+ synapses through several potential mechanisms.

At the glutamatergic Calyx of Held, α2δs set synaptic gain by controlling the quantity of synaptic VGCCs (Milanick et al., 2025). Even after deletion of all presynaptic α2δ isoforms at the Calyx, VGCCs are normally clustered and positioned relative to vesicle release sites, as defined by RIM1/2 or Munc13 co-localization by immunoelectron microscopy (Milanick et al., 2025). Although this result appears at odds with our calcium chelator results, active zones are organized differently at different synapses (Cano and Tabares, 2016), and it is possible that α2δs could contribute to subsynaptic localization and nanodomain coupling at hippocampal PV+ synapses but not at the Calyces of Held, which possess a unique anatomical organization.

Alternatively, even though RIMs and Munc13 are critically involved in vesicle priming and release (Betz et al., 1998; Han et al., 2011), the co-localization required for functional calcium-release coupling may involve other Ca^2+^-binding proteins in PV+ terminals. Unfortunately, hippocampal PV+ interneuron terminals are poorly amenable to analysis of synaptic ultrastructure, and confocal microscopy is limited in its subsynaptic resolving capacity. Hopefully future work will be able to more easily quantify presynaptic VGCC concentration and sub-terminal localization at these terminals, and to identify whether α2δ proteins play a role in the localization of VGCCs relative to putative critical presynaptic calcium sensors such as the synaptotagmins.

Together, current and prior data all highlight the various roles of α2δ proteins in neuronal function. Beyond VGCC trafficking, functional coupling to effector mechanisms, and synaptogenesis, these roles can even include regulation of potassium conductances through interactions with BK channels, transsynaptic receptor recruitment, or even activity-dependent extracellular α2δ release to affect heterosynaptic receptor recruitment (Zhang et al., 2018; Geisler et al., 2019; Dos Santos et al., 2026). Future work will hopefully delineate which of these many functions α2δ proteins perform at specific synapses, and whether they are cell-type, target, and/or α2δ isoform-specific.

### Dysfunctional synaptic inhibition by parvalbumin-expressing interneurons may underlie hyperexcitability and epilepsy in α2δ mutant mice

Our data provide insights into how α2δ-2 loss specifically contributes to PV+ interneuron dysfunction. PV+ interneurons play a critical role in hippocampal circuit modulation (Zhang et al., 2025), and disrupted PV+ interneuron function has been linked to epileptic phenotypes (Zhang and Buckmaster, 2009). As the *α2δ-2* KO mouse is epileptic and exhibits increased dentate gyrus activation *in vivo* (Barclay et al., 2001; Danis et al., 2024), we propose that decreased PV-mediated inhibition underlies these phenotypes.

α2δ-2 is selectively expressed in PV+ interneurons across the brain (Cole et al., 2005), and thus a similar GABA release phenotype might exist in any PV+ neurons that selectively express the α2δ-2 isoform. The established links between the dentate gyrus and seizure emergence (Zhang and Buckmaster, 2009; Krook-Magnuson et al., 2015) suggest that *α2δ-2* loss leading to dentate PV+ interneuron dysfunction could be a substantial component.

Seizures in *α2δ-2* null mice are similar to those observed in absence epilepsy, which typically is associated with thalamocortical dysfunction (Barclay et al., 2001; Crunelli et al., 2020). As α2δ-2 is expressed in cortical PV+ cells (Cole et al., 2005), dysfunction in cortical PV+ neurons might contribute to epilepsy in α2δ-2-deficient animals, if cortical PV+ neurons share the same GABA release phenotypes. This would complement recent evidence suggesting that cortical PV+ neuron dysfunction plays a central role in the pathogenesis of absence epilepsy, and that this may specifically relate to P/Q-type VGCC dysfunction in PV+ cells, which are highly dependent on this channel subunit for GABA release (Hefft and Jonas, 2005; Miao et al., 2026; Song et al., 2026).

Identification of a critical role for α2δ-2 in mediating functional coupling in PV+ terminals contributes to a growing understanding of the roles of α2δ proteins in the brain, and how a specific genetic mutation which disrupts synaptic VGCC signaling at synapses might ultimately underlie the epileptic phenotypes in *α2δ-2* knockout mice and humans with α2δ-2-related genetic epilepsy.

## Acknowledgements

We thank Gary Westbrook and his lab members, Melissa Hermann, Cory Butler, and Skyler Jackman for helpful discussions of our data, Drs. Sergey Ivanov and Lino Tessarollo for generously providing mutant mice, the OHSU Advanced Light Microscopy Core (RRID:SCR_009961) and Jennifer Jahncke for assistance with image analysis, Alyssa Danis and Arielle Isakharov for technical assistance, and Milana Krush and members of the Schnell lab for feedback and support.

